# Biochemical and structural characterization of glycosyltransferase family 61 proteins reveal a key determinant of sugar donor specificity

**DOI:** 10.1101/2025.07.18.665579

**Authors:** Hsin-Tzu Wang, Jose Henrique Pereira, Andy M. DeGiovanni, Pradeep Kumar Prabhakar, Digantkumar Chapla, Kelley W. Moremen, Breeanna R. Urbanowicz, Paul D. Adams, Henrik V. Scheller

## Abstract

Xylan, the most abundant non-cellulosic polymer in plant cell walls, is structurally diverse, especially in grasses where it is heavily substituted with arabinofuranose and further modified by various residues. Common substitutions across species include glucuronic and 4-*O*-methyl-glucuronic acid. Arabinose and xylose sidechains are synthesized by glycosyltransferase family 61 (GT61) proteins, many of which remain uncharacterized in plants, with limited structural and mechanistic understanding. In this study, we identified two novel GT61 enzymes in *Sorghum bicolor*, functioning as xylan arabinosyltransferase (SbXAT) and xylan xylosyltransferase (SbXXT). We resolved the crystal structure of SbXAT, which exhibits a GT-B fold with two Rossmann-like domains linked by a cleft that accommodates the catalytic site. Structural comparison with a predicted SbXXT model revealed a substrate-binding residue critical for sugar donor specificity, validated through site-directed mutagenesis and enzymatic assays. These findings enhance understanding of xylan biosynthesis and provide a foundation for engineering glycosyltransferases and predicting their functions.

## Introduction

Xylans are the most abundant non-cellulosic polysaccharides in biomass, and they are ubiquitous in plant cell walls of different species with highly diverse structures ^1^. In grasses, xylans are composed of backbones formed by β-1,4-linked D-xylopyranose (Xyl*p*) residues that are frequently substituted with α-L-arabinofuranose (Ara*f*) residues at *O*-2 and/or *O*-3 positions ^2,3^. Besides arabinosyl substitutions, xylan backbones in grasses are often acetylated at *O*-2 and/or *O*-3 and modified by α-1,2-linked D-glucuronic acid (Glc*p*A) and 4-*O*-methyl-D-glucuronic acid (MeGlc*p*A) to a lesser extent ^4^. A distinct feature found in grass glucuronoarabinoxylan (GAX) is that the α-1,3-linked Ara*f* side chains can be further esterified by 5-*O*-*p*-coumaroyl or 5-*O*-feruloyl units, where the latter can cross-link with another GAX or lignin, which can strengthen the cell wall structure and increase recalcitrance ^5^. The 5-*O*-feruloyl-L-arabinosyl side chains frequently contain a β-2-*O*-Xyl*p* ^6,7^, and these Xyl*p*-Ara*f* substituents can be further extended with β-1,4-galactose (Gal*p*) residues ^8^. Finally, the presence of arabinobiose substituents formed by adding an additional α-1,2-Ara*f* to α-1,3-Ara*f* side chains has also been reported ^9^.

In plants, several members of the glycosyltransferase family 61 (GT61) have been found to be involved in addition of pentose substituents of GAX, and have been divided into three clades based on phylogenetic analysis ^10^ (Fig. S1). Clade A has been extensively expanded in grasses and contains xylan arabinosyltransferases (XATs) and xylan xylosyltransferases (XXTs) involved in GAX substitution in grasses, while no dicot GT61 has been functionally characterized so far in Clade A. Clade B has a relatively even amount of dicot and monocot members with a few biochemically characterized members in rice ^11,12^ and Arabidopsis, where MUCI21 has been suggested to add β-1,2-Xyl*p* directly to the xylan backbone in seed mucilage ^13^. Clade C GT61s are involved in protein *N*-glycosylation.

The first evidence of GT61 involvement in xylan substitution was the observation of decreased α-1,3-arabinosylated xylans in wheat *TaXAT1* RNAi transgenic endosperm. A gain-of-function study of heterologously expressed rice *OsXAT2*, *OsXAT3*, and wheat *TaXAT2* in Arabidopsis resulted in arabinosylation of xylan, which is not a typical modification in Arabidopsis, indicating that those wheat and rice GT61s possess XAT activity ^10^. Later, OsXAT2, OsXAT3 and their homologs in rice and other grass species were biochemically characterized as XATs through *in vitro* activity assays ^11^.

A distinct xylosyltransferase function has been proposed in a study of the rice *xylosyl arabinosyl substitution of xylan 1* (*xax1*) mutant, which showed the absence of β-1,2-Xyl*p*-α-1,3-Ara*f* side chains on xylan ^14^. However, an alternate function of XAX1 in transferring hydroxycinnamic acid-modified arabinosyl substitutions to xylans was later proposed, based on the decreased level of ferulated Ara*f* and diferulated side chains regardless of their xylosyl substitutions in the *xax1* mutant ^15^. Meanwhile, rice OsXAXT1, together with its homologs from *Brachypodium*, switchgrass, and maize were heterologously co-expressed with OsXAT2 in the Arabidopsis *gux1/2/3* mutant that lacks glucuronic acid substitutions, and the results showed the presence of β-1,2-Xyl*p*-α-1,3-Ara*f* disaccharide side chains on xylans in the transgenic lines, indicating a conserved function of xylosylating α-1,3-Ara*f* residues on xylans by GT61s in grass species ^16^. Besides transferring Xyl*p* onto α-1,3-Ara*f* side chains, a direct transfer of Xyl*p* to the xylan backbone by GT61s has also been reported in three rice GT61s (OsXYXT1 in Clade A, OsXXAT1 and OsXYXT2 in Clade B) through *in vivo* (OsXYXT1 only) and *in vitro* analysis ^12,17^. Interestingly, OsXXAT1 has been found to exhibit dual functions as it also showed XAT activity although with a relatively lower level compared to its XXT activity ^12^.

Here, we biochemically characterized two sorghum GT61s as a xylan arabinosyltransferase (SbXAT) and a xylan xylosyltransferase (SbXXT) residing in Clade A and Clade B, respectively. The crystal structure of SbXAT reveals a key determinant for UDP-sugar specificity in the catalytic pocket based on molecular docking and validation through site-directed mutagenesis. The biochemical and structural results presented here broaden our understanding of xylan biosynthesis, which can be further applied to protein engineering and plant cell wall modification to generate better feedstocks for agriculture and biofuel production.

## Results

### Identification and biochemical characterization of sorghum GT61 enzymes

To identify the glycosyltransferase activity of the GT61 proteins, we expressed the selected proteins in human embryonic kidney cells (HEK cells). In order to evaluate NDP-sugar donor specificity, we evaluated the ability of the enzymes to transfer a sugar from different donor substrates (e.g. UDP-arabinofuranose, UDP-xylose, UDP-glucose, UDP-glucuronic acid, UDP-galactose, UDP-galacturonic acid) onto a xylohexaose (Xyl_6_) acceptor. Analysis of the reaction products by matrix-assisted laser desorption/ionization time-of-flight mass spectrometry (MALDI-TOF MS) showed that Sb3G094600 (SbXAT) can only utilize UDP-Ara*f* as the sugar donor to catalyze arabinosylation on Xyl_6_, while Sb2G401600 (SbXXT) selectively uses UDP-Xyl and transfers xyloses to Xyl_6_ (Fig. 1). Notably, SbXAT was able to generate highly arabinosylated products in the first three hours of the reaction (Fig. S2a), and could attach up to five arabinoses to the Xyl_6_ acceptor after overnight incubation (Fig. 1a); while the xylosylated products generated by SbXXT could only be detected after an overnight reaction, and only two xyloses were transferred to Xyl_6_ (Fig. 1b, Fig. S2b).

**Fig. 1.**
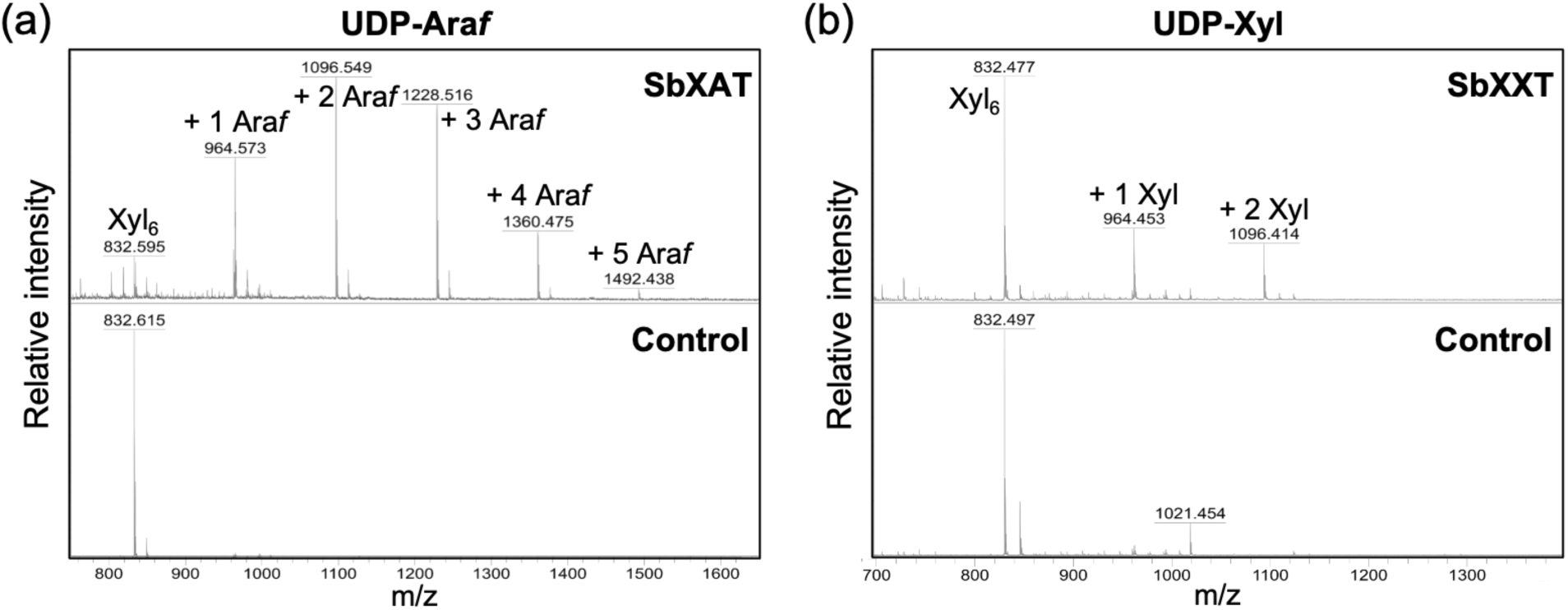
MALDI-TOF MS analysis of the products generated by SbXAT (a) and SbXXT(b) using UDP-Ara*f* and UDP-Xyl, respectively, as the donor substrates. Xyl_6_ was used as the acceptor in the *in vitro* assays and the products were analyzed after overnight incubation. Each transfer of an Ara*f* or Xyl increases the mass of the Xyl_6_ acceptor by 132 Da as shown by annotated [M+Na]^+^ ions. Analysis after 2 and 3 h incubation is shown in Fig. S2.

After verifying the donor specificities of the GT61 proteins, we further investigated their acceptor specificities using different oligosaccharides (e.g. xylobiose (Xyl_2_), xylotriose (Xyl_3_), xylotetraose (Xyl_4_), xylopentaose (Xyl_5_), arabinosylated xylotetraose, xyloglucoheptasaccharide, mixed-linkage glucotetraose, and mannohexaose) in the activity assays. The results showed that both GT61s are xylan-specific and SbXXT only uses unbranched xylo-oligosaccharides as the acceptor (Fig. 2). Among the linear xylo-oligosaccharides with different degrees of polymerization (DP), Xyl_6_ was the preferred substrate for both SbXAT and SbXXT, and the lowest DP that can be utilized by the enzymes are DP3 (e.g. Xyl_3_) and DP4 (e.g. Xyl_4_) for SbXAT and SbXXT, respectively.

**Fig. 2.**
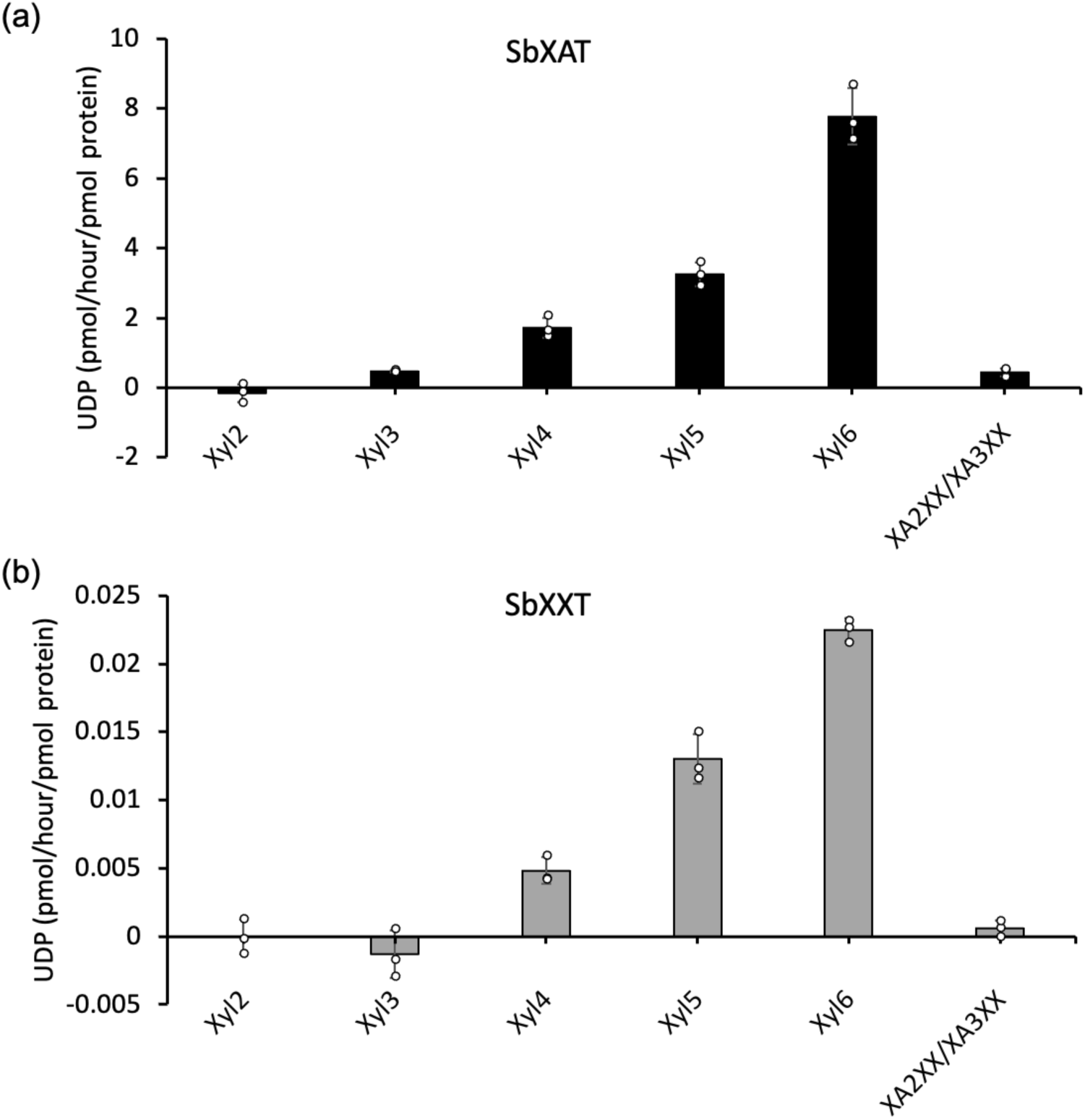
Acceptor substrate specificity analysis of SbXAT (a) and SbXXT (b). Activity with different xylo-oligosaccharides was determined by quantification of UDP. UDP-Ara*f* was used as the donor substrate for SbXAT, while UDP-Xyl was used for SbXXT. Data are represented as means ± SD from three independent reactions (n = 3). Xyl_2_, xylobiose; Xyl_3_, xylotriose; Xyl_4_, xylotetraose; Xyl_5_, xylopentaose; Xyl_6_, xylohexaose; XA^2^XX/XA^3^XX, mixture of arabinosylated xylotetraoses with Ara*f* attached at *O*-2 or *O*-3 positions of the second Xyl residue from the terminal end.

### Crystal structure of the xylan arabinosyltransferase SbXAT

The SbXAT protein sequence is composed of 685 amino acid residues, with residues 35 to 53 predicted as a transmembrane domain. Analysis of the AlphaFold ^18^ model of SbXAT showed a large disordered N-terminal region (residues 1-293) with the catalytic domain comprising residues 294 to 685. Since the AlphaFold model showed a potential α-helix starting at position 205, we created a truncated SbXAT (amino acids 198-685) construct for crystallography studies.

The crystal structure of SbXAT was solved at 2.0 Å resolution (Table S1). The first residue visible from the electron density map was Trp294, indicating that residues 198-293 are disordered in the crystal structure, in close agreement with the AlphaFold model disorder prediction. The entire SbXAT catalytic domain (residues 294-685) was clearly built on the electron density map with the exception of the last 11 C-terminal residues. The general structure of SbXAT adopts a GT-B fold ^19^, which is characterized by two Rossmann-like domains (N-lobe and C-lobe) that are positioned in close proximity creating a cleft that locates the SbXAT active site (Fig. 3a). The N-lobe comprises residues 294-493 and the C-lobe comprises residues 508-685. There is a high structural conservation between members of the GT-B family, especially the C-lobe domain. Structural changes are more evident in the N-lobe domains, in the regions pointing towards the active site, which have evolved to accommodate different acceptors ^20^. GT-B fold members like SbXAT have an inverting glycosylation catalytic mechanism ^21^ where the catalytic base is provided by an active-site side chain (such as Asp, Glu or His). In SbXAT the catalytic residue is His397 and/or His564 (Fig. 3b). One glycosylation site at residue Asn321 was observed in the crystal structure of SbXAT (Fig. S3).

**Fig. 3.**
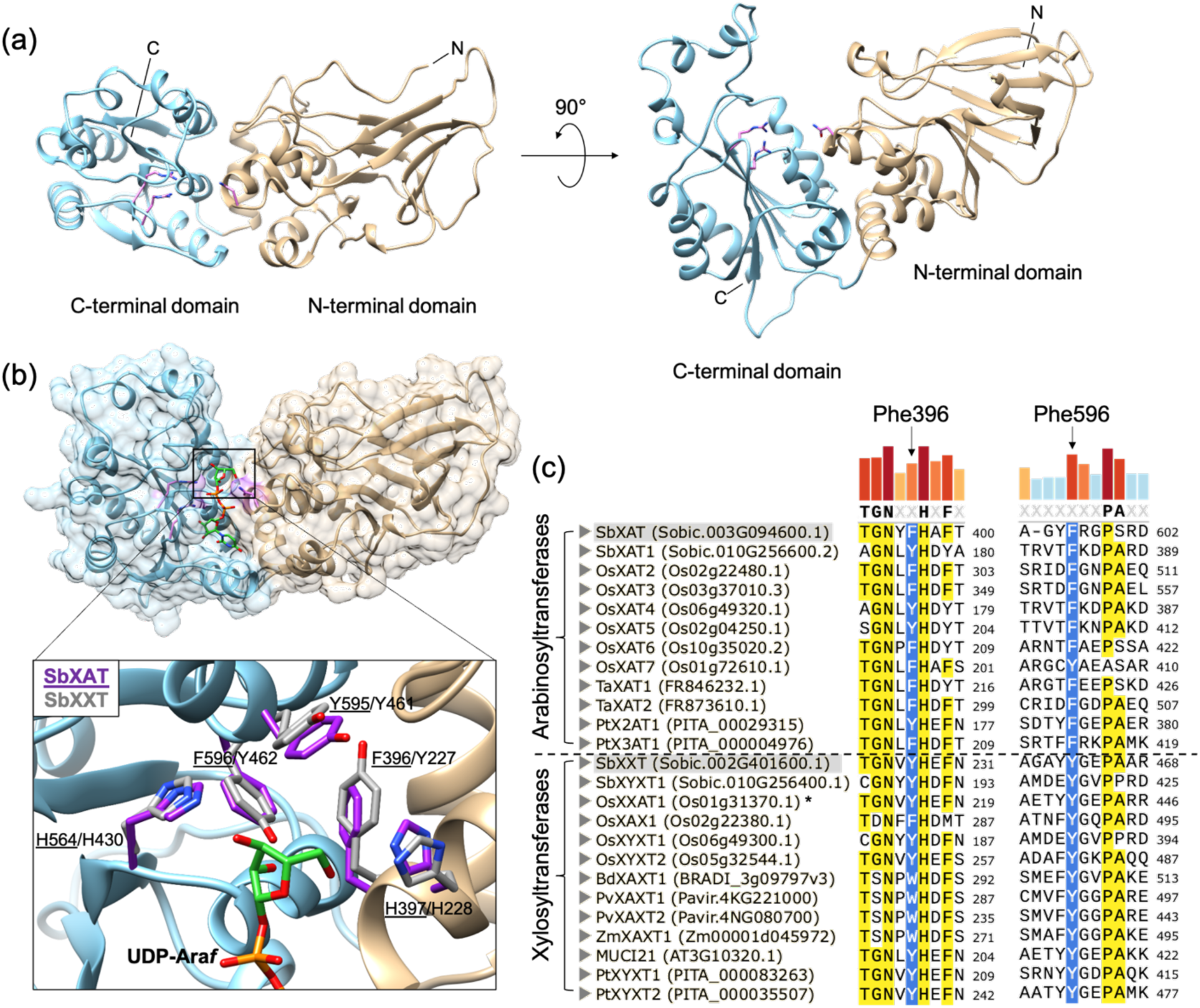
Crystal structure of SbXAT and molecular docking of UDP-Ara*f* to identify the donor binding site. **a**, The overall crystal structure of SbXAT is shown in ribbons with the N-terminal domain (tan) and C-terminal domain (blue) highlighted. The side chains of the three residues (Arg515, Arg519, Asn394) hypothetically interacting with the diphosphate moiety of the UDP-sugar according to the POMGNT2 co-complex (PDB: 7E9J) are shown as sticks (magenta). **b**, Structure of the SbXAT docked model is shown in ribbons with a surface (tan, NTD; blue, CTD; magenta, substrate-binding residues), and the docked UDP-Ara*f* in the binding cleft is shown in sticks (green). Inset: zoom-in view of the substrate-binding site of SbXAT that shows the five ring amino acids (purple; underlined annotations) interacting with the sugar moiety of the UDP-Ara*f*, the aligned amino acids from the predicted structure of SbXXT are colored in gray with annotations. **c**, Protein sequence alignment of the characterized xylan-specific GT61s. The equivalent residues in different arabinosyltransferases/xylosyltransferases to Phe396 and Phe596 in SbXAT are highlighted in blue. The two GT61s identified in our study are shadowed. Colored bars on the top represent the consensus sequence, and residues with > 70% consensus threshold are highlighted in yellow. MUSCLE was used for multiple sequence alignment. *, OsXXAT1 also shows arabinosyltransferase activity with a lower level.

### Protein engineering of GT61 reveals a key residue involved in sugar donor recognition

To understand the enzyme-substrate interaction of the GT61 proteins, we carried out molecular docking to dock UDP-Ara*f* to the active site of the SbXAT crystal structure (Fig. 3b). Superposition of the SbXAT structure against the glycosyltransferase POMGNT2 structure in complex with UDP ^22^ showed a similar binding position of UDP compared to the docking results (Fig. S4a). Moreover, all the three residues of POMGNT2 identified as important for interacting with the α- and β-phosphates of UDP (Asn163, Arg294 and Arg298) are fully conserved in SbXAT (Asn394, Arg515 and Arg519) indicating similar mode of recognition of the NDP-sugar donor between the two proteins (Fig. S4b). Mutagenesis experiments replacing any of these residues by alanine completely abolished the enzymatic activity of POMGNT2 ^22^.

The docked model showed that there are 3 aromatic residues (Phe396, Tyr595, and Phe596) and 2 histidines (His397 and His564) in the catalytic site of SbXAT that possibly interact with the sugar moiety of UDP-Ara*f* through hydrogen bonding and hydrophobic interactions. To further evaluate the individual importance of those ring amino acids in recognizing specific sugar moieties, we aligned the crystal structure of SbXAT with the predicted model of SbXXT acquired from AlphaFold Protein Structure Database (https://alphafold.ebi.ac.uk/) ^18,23^ and compared the amino acids in the equivalent positions. The two histidine residues in the binding pocket of SbXXT are conserved, but instead of having phenylalanines in the position of Phe396 and Phe596 in SbXAT, SbXXT harbors tyrosines in the equivalent positions (Tyr227 and Tyr462) (Fig. 3b). To investigate the equivalent amino acid residues in all known xylan arabinosyltransferases and xylosyltransferases in GT61, we sequence aligned the 25 characterized xylan-specific GT61s from the CAZy (Carbohydrate Active Enzyme) database (https://www.cazy.org, April 2025 ^24^) and GenBank® (https://www.ncbi.nlm.nih.gov/genbank/, April 2025 ^25^). The results showed that the xylan-specific arabinosyltransferases (11 out of 12, with an exception of OsXAT7 having a tyrosine) preferentially have a phenylalanine in the equivalent position of residue Phe596 in SbXAT, while all the xylan xylosyltransferases have tyrosines instead (Fig. 3c). No preferential amino acids in the equivalent position of Phe396 in SbXAT could be observed among the characterized xylan-specific GT61s (Fig. 3c).

Having either a phenylalanine or a tyrosine specifically occupied in the conserved position in different functional GT61s strongly suggests a key role that these residues play in recognizing UDP-sugars and thus determine the enzymes’ functionalities. To evaluate if phenylalanines and tyrosines are key determinants for the donor substrate specificity of GT61s, we carried out site-directed mutagenesis substituting Phe596 in SbXAT with a tyrosine (SbXAT F596Y), and substituting the equivalent residue Tyr462 in SbXXT with a phenylalanine (SbXXT Y462F). The *in vitro* activity assays followed by MALDI-TOF MS analysis showed that the native arabinosyltransferase activity of SbXAT was negatively affected in the F596Y variant since less arabinosylated products were detected from the overnight reaction (Fig. 4a). Meanwhile, SbXAT F596Y was able to utilize UDP-xylose as the donor substrate and transferred up to two xyloses onto Xyl_6_, indicating that F596Y mutation tunes the substrate specificity of SbXAT and turns the enzyme into a bi-functional xylan arabinosyltransferase/xylosyltransferase. Conversely, the assays of SbXXT Y462F showed similar results that the original xylosyltransferase activity dropped while arabinosyltransferase activity was gained, based on less amount of xylosylated Xyl_6_ produced by SbXXT Y462F, and a gained arabinosyltransferase activity based on the presence of the arabinosylated Xyl_6_ (Fig. 4b).

**Fig. 4.**
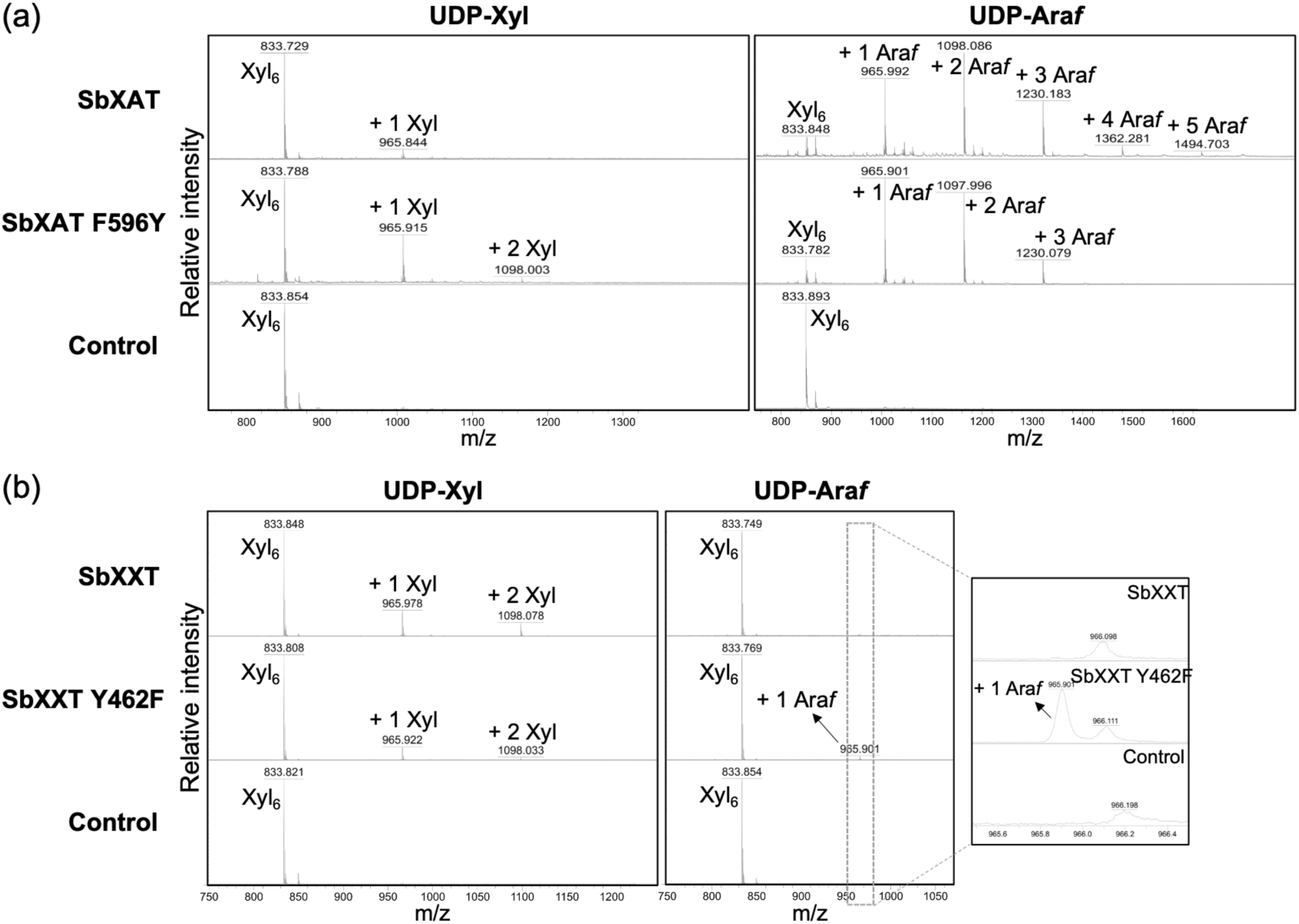
Modified donor substrate selectivity of GT61 variants. MALDI-TOF MS analysis of the products generated using UDP-Ara*f* or UDP-Xyl as donor substrates and Xyl_6_ as the acceptor in overnight *in vitro* assays. **a,** SbXAT and SbXAT F596Y. **b,** SbXXT and SbXXT Y462F; the inset is the zoomed-in spectra of the m/z range 965.5 – 966.5. Each transfer of an Ara*f* or Xyl increases the mass of the Xyl_6_ acceptor by 132 Da as shown by annotated [M+Na]^+^ ions. Control assays were incubated without enzyme.

To further investigate the properties of the GT61 variants, we evaluated the binding affinities of GT61 and their variants for different UDP-sugar substrates through microscale thermophoresis (MST). The dissociation constant (*K*_d_) of SbXAT for its native donor substrate, UDP-Ara*f*, was 244 μM, while the *K*_d_ of SbXAT F596Y for UDP-Ara*f* was 531 μM (Fig. 5a, left), demonstrating a weaker interaction between the variant and UDP-Ara*f*, consistent with the lower arabinosyltransferase activity of SbXAT F596Y observed in the *in vitro* assays (Fig. 4a). Conversely, the *K*_d_ of SbXAT F596Y for UDP-Xyl was 221 μM, which is lower than that of SbXAT (373 μM) (Fig. 5a, right), indicating a better binding affinity of the variant for UDP-Xyl and supporting its gained catalytic ability of xylosylating Xyl_6_ as shown by the activity assays (Fig. 4a). Likewise, SbXXT Y462F showed a lower *K*_d_ for UDP-Ara*f* (19 μM) compared to the corresponding wildtype enzyme (*K*_d_ = 26 μM), although with a large standard deviation. It also showed a higher *K*_d_ for UDP-Xyl (15 μM) compared to SbXXT (*K*_d_ = 13 μM) (Fig. 5b), demonstrating that the Y462F mutation weakened the binding affinity between SbXXT and its native donor substrate, UDP-Xyl. Lastly, we measured the *K*_d_ of GT61 for the acceptor Xyl_6_, and the results showed that binding only occurred in the presence of UDP, and revealed SbXXT (*K*_d_ = 22 μM) displays a higher binding affinity for Xyl_6_ relative to SbXAT (*K*_d_ = 178 μM) (Fig. S5).

**Fig. 5.**
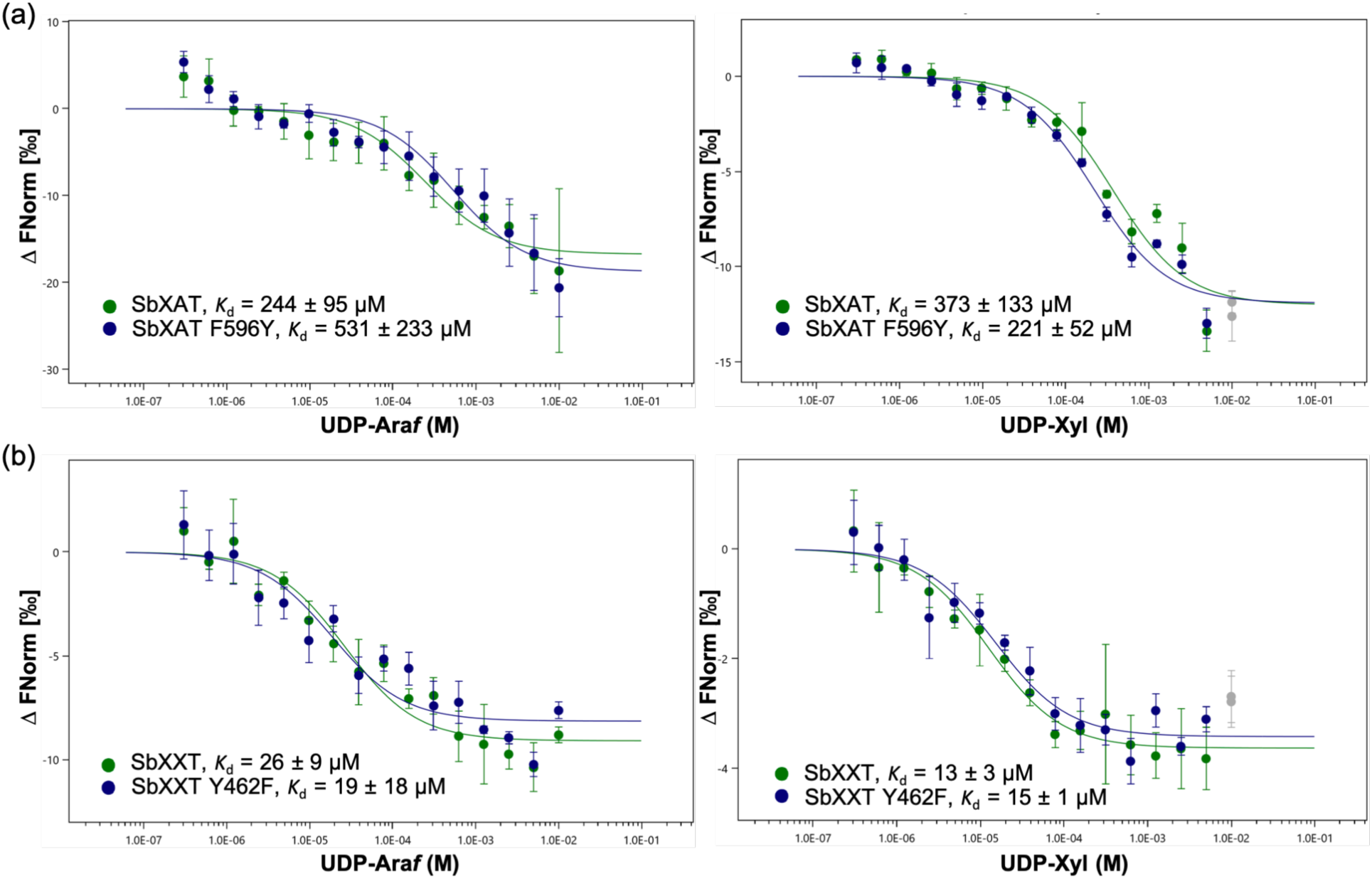
Substrate-binding affinity analysis of SbXAT, SbXXT, and their variants for UDP-Ara*f* and UDP-Xyl through microscale thermophoresis (MST). Normalized binding curves of SbXAT and SbXAT F596Y (**a**), and SbXXT and SbXXT Y462F (**b**), for UDP-Ara*f* (left) and UDP-Xyl (right) that yield the dissociation constants (*K*_d_) ± SD as indicated in the figure. Concentration of enzymes used in each assay was kept constant (200 nM) while the UDP-sugar concentrations varied from 0 to 10 mM. Green, wildtype GT61 proteins; blue, GT61 variants; gray, ignored data points due to protein aggregation. Error bars represent means ± SD from three technical replicates (n = 3).

We next investigated how point mutations affect GT61 enzyme activity, focusing on substrate specificity and catalysis with or without the acceptor Xyl_6_. Incubation of SbXAT and its F596Y variant with UDP-Ara*f* revealed that both enzymes showed greater activity in the presence of Xyl_6_, as evidenced by increased UDP production (Fig. 6a). This confirms that both possess arabinosyltransferase activity toward Xyl_6_. Notably, SbXAT and SbXAT F596Y also hydrolyzed UDP-Ara*f* in the absence of acceptor, though SbXAT consistently showed higher activity, indicating that Phe596 is important for both glycosyltransferase and hydrolase functions. In contrast, neither SbXXT nor its Y462F variant exhibited activity with UDP-Ara*f*, regardless of Xyl_6_. However, when using UDP-Xyl, SbXAT F596Y demonstrated significantly enhanced activity — nearly 5-fold higher than wildtype SbXAT in the presence of Xyl_6_ and also showed increased hydrolysis in the absence of the acceptor. This suggests that the F596Y mutation broadens substrate specificity and improves catalytic activity with UDP-Xyl. Meanwhile, SbXXT and its Y462F variant were only active with UDP-Xyl in the presence of Xyl_6_, with reduced activity observed in Y462F, highlighting the critical role of Tyr462 in catalysis. Minimal hydrolytic activity in the absence of acceptor suggests these enzymes require Xyl_6_ for activity.

**Fig. 6.**
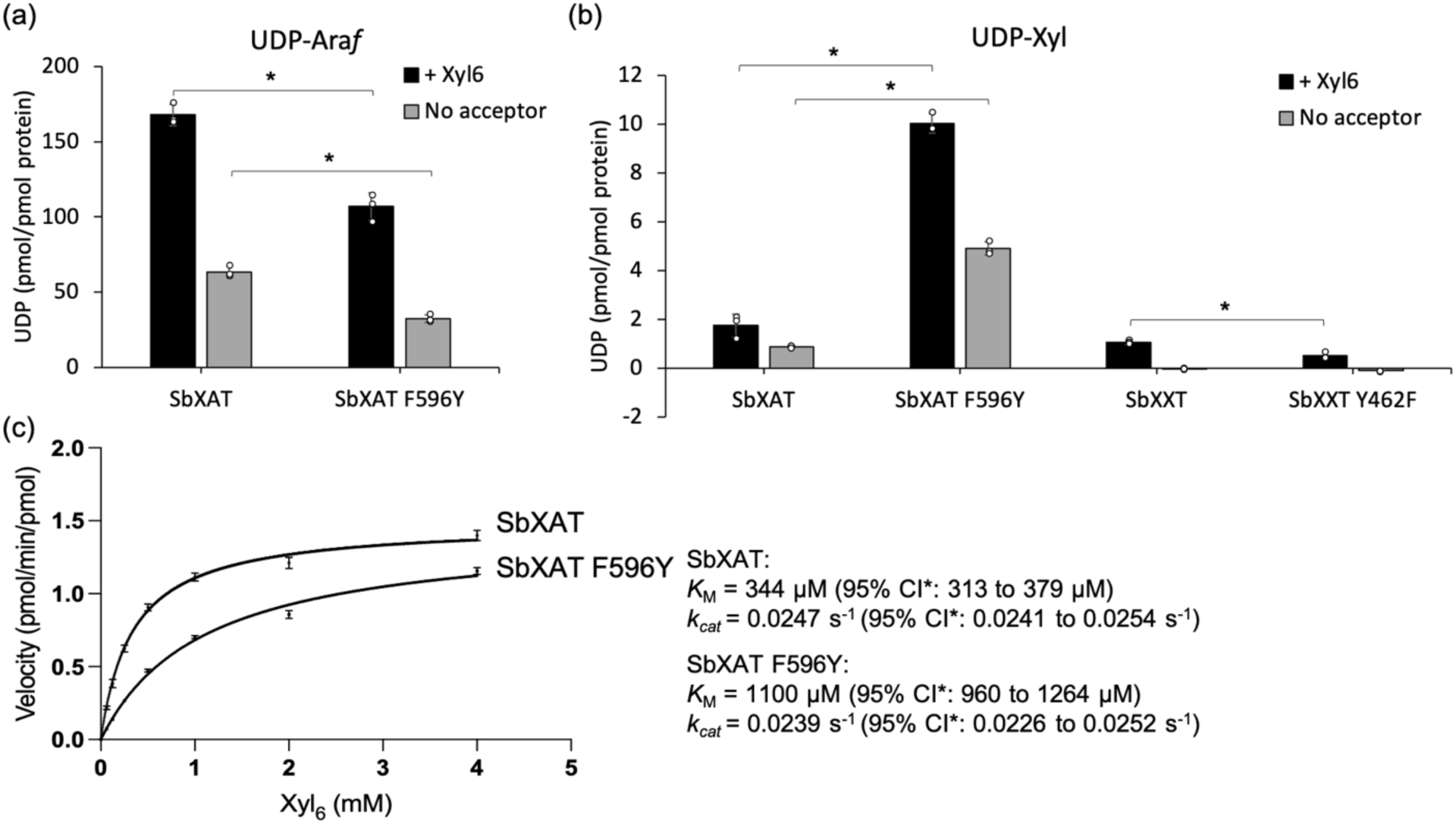
Enzymatic activities of GT61 enzymes and their variants. **a**, UDP-Ara*f* catalysis by SbXAT and SbXAT F596Y after overnight incubation with and without Xyl_6_. **b,** UDP-Xyl catalysis by SbXAT, SbXAT F596Y, SbXXT, and SbXXT Y462F after overnight incubation with and without Xyl_6_. UDP by-product generated during the reactions was quantified by the UDP-Glo™ assay kit. Statistical significance of each variant (SbXAT F596Y or SbXXT Y462F), compared to the corresponding wild-type enzyme (SbXAT or SbXXT, respectively), was determined by the two-tailed Student’s *t*-test. *, *P* < 0.01. **c**, Michaelis-Menten curves and kinetic parameters of SbXAT and SbXAT F596Y for Xyl_6_ when using UDP-Ara*f* as the donor substrate. Confidence intervals (CI) of *K*_M_ and *k_cat_* were calculated at 95% confidence level. Error bars represent means ± SD from three replicates of independent reactions (n = 3).

Kinetic analysis using the UDP-Glo™ assay with fixed UDP-Ara*f* and varied Xyl_6_ concentrations revealed a higher *K*_M_ for SbXAT F596Y (1100 μM) compared to SbXAT (344 μM), with similar turnover numbers (*k_cat_* ≈ 0.024 s⁻¹). Consequently, the catalytic efficiency (*k_cat_*/*K*_M_) of SbXAT F596Y was significantly lower (21.7 M⁻¹s⁻¹) than that of wildtype SbXAT (71.8 M⁻¹s⁻¹), suggesting reduced affinity for Xyl_6_ in the mutant. These results provide mechanistic insights into how specific GT61 residues modulate sugar donor preference and catalytic performance. Taken together, swapping the equivalent phenylalanine and tyrosine in the binding sites of SbXAT and SbXXT resulted in gained xylosyltransferase and arabinosyltransferase activities, respectively, and their native glycosyltransferase activities have been weakened by the point mutation, suggesting a key role that the phenylalanine/tyrosine in close proximity to the sugar moiety play in recognizing and interacting with the UDP-sugar, and it demonstrates that they are key determinants of donor substrate specificity of GT61s.

## Discussion

GT61 family is widely expanded in grass species, and several members have been functionally characterized as xylan arabinosyltransferases (XATs), xylosyltransferases (XXTs), or xylosyl/arabinosyltransferases (XXATs) that possesses dual activities involved in xylan substitution ^10–12,14,16,17^. In sorghum, there are 29 members in the GT61 family, but only two of them have been biochemically characterized ^11^. In order to investigate the potential function possessed by the expanded GT61 proteins, we expressed five protein candidates residing in the less explored subclades in clade A and B in the GT61 phylogenetic tree and biochemically characterized two of them, SbXAT and SbXXT. Interestingly, the two GT61s showed distinct features, with SbXAT being highly active and able to transfer up to five arabinoses onto the Xyl_6_ acceptor after an overnight reaction, while SbXXT is a slow enzyme that is only able to generate a low abundance of xylosylated products with a maximum of two xyloses attached to Xyl_6_ (Fig. 1). The distinct enzymatic efficiencies presented by SbXAT and SbXXT might reflect the natural feature of xylan structures in grass species, where xylan backbones are highly substituted by Ara*f* ^1^, while xylosyl substituents directly attached to xylan backbones have not been reported in grass cell walls, and have only been observed in dicot tissues like psyllium husk ^26,27^ and Arabidopsis seed mucilage ^13^. These unique xylan structures have been suggested to be essential for pectin attachment to the seed surface based on the *muci21* mutant study in Arabidopsis. MUCI21 is required to facilitate the addition of β-1,2-xyloses to the xylan backbone although the transferase activity has not been biochemically confirmed ^13^. The biochemical feature of SbXXT shown in our study together with the previously identified xylosyltransferases in various grass species ^11,12,17^ suggests that xylosylated xylans could be potentially present in the cell walls of commelinid monocots, although they have not been identified, probably due to low abundance.

The structure of the mammalian GT61 enzyme POMGNT2 includes an N-terminal catalytic domain with two Rossmann-like folds (N-lobe and C-lobe) and a C-terminal fibronectin type III (FnIII) domain, which mediates dimer formation via interaction with the N-lobe of another protomer ^22,28^. Key residues interacting with UDP and the acceptor peptide have been identified, supporting an ordered sequential bi-bi catalytic mechanism. ^22^. Here, we present the first structure of a plant GT61 protein, SbXAT, which lacks an FnIII domain and exhibits a typical GT-B fold comprising two Rossmann-like domains: an N-terminal domain (NTD) for acceptor binding and a C-terminal domain (CTD) for donor binding ^20^.

SbXAT’s NTD features a conserved NxxHxxxD motif with His397 being equivalent to the proposed catalytic base His166 of bovine POMGNT2. Additionally, His564 of the HGAxxT motif in the CTD, near the UDP-Ara*f* sugar moiety, corresponds to His345 in human POMGNT2, also proposed as a catalytic base. Asp401 (from NxxHxxxD) may stabilize the catalytic His397, consistent with findings from a GT-B fold plant enzyme from the GT1 family, UGT71G1 ^29^. Asn394, Arg515, and Arg519 (from NxxHxxxD and RxxxRxxxN motifs) align with critical residues in POMGNT2 that play a key role in stabilizing UDP-sugars ^22^.

The catalytic and substate-binding residues mentioned above are highlighted in an amino acid sequence alignment of GT61 proteins with indicated conserved motifs (Fig. S6). Lastly, a flexible N-terminal loop, observed in various Golgi-localized plant glycosyltransferases involved in cell wall biosynthesis ^30^, was also present in our SbXAT protein construct. Notably, a flexible loop with unclear function connecting the N-terminal transmembrane domain and the catalytic domain has been observed in different Golgi-localized enzymes involved in plant cell wall biosynthesis ^30^. Though not resolved in the crystal structure due to low electron density, including a larger part of this region in the protein construct enhanced expression and enzymatic activity (Fig. S7). The identification of these conserved structural features in SbXAT provides important insights into GT61 catalytic mechanisms and substrate interactions in plants.

Aromatic side chains are frequently enriched in the substrate-binding pocket of carbohydrate-processing enzymes, where electron-rich aromatic rings interact with the electropositive carbohydrate C-H bonds via CH-π interactions that contribute to protein recognition of glycans ^31–34^. Other interactions between aromatic amino acids and carbohydrate ligands such as hydrogen bonding are also involved in binding ^32^. The potential UDP-sugar-binding residues shown in the catalytic pocket of our SbXAT docked model also include five aromatic amino acids and histidines in close proximity with the sugar moiety of UDP-Ara*f* (Fig. 3b). Through structural comparison between SbXAT and the predicted model of SbXXT, we found two of the five residues in SbXAT (Phe396 and Phe596) having different side chains from their equivalent amino acids in SbXXT (Tyr227 and Tyr462, respectively). Sequence alignment of the characterized GT61s further indicates that only the C-terminus-residing phenylalanine (i.e. Phe596 in SbXAT) dominates the equivalent position among xylan arabinosyltransferases from various species, while tyrosine occupies the same position in all characterized xylan xylosyltransferases (i.e. Tyr462 in SbXXT). This observation suggested a crucial role of phenylalanine and tyrosine in the CTD in determining donor substrate specificity, which was further validated by our site-directed mutagenesis results showing gained glycosyltransferase activities in SbXAT F596Y and SbXXT Y462F variants.

The CTDs are highly conserved among UDP-glycosyltransferases (UGTs) that transfer sugars from UDP-sugars to various acceptor molecules, which is most likely due to the chemical and structural similarity of sugar donors they recognize ^35^. Several studies on plant secondary metabolite glycosyltransferases from GT1 show that site-directed mutagenesis of the residues residing in the CTD resulted in a shifted sugar donor specificity ^36–39^. These enzymes have a highly conserved Plant Secondary Product Glycosyltransferase (PSPG) motif consisting of 44 amino acid residues, and structural studies show that it interacts with UDP-sugars through its highly conserved residues, with residues at the 43rd and 44th position crucial to the selectivity of UGTs for different sugar donors ^40^. The PSPG motif is not present in the plant GT61 family, but the HGAxxT conserved motif mentioned above occupies the same position that the PSPG motif does in plant secondary metabolite UGTs.

Although the NTD in UGTs primarily interacts with acceptor molecules, there are NTD-residing residues also critical for donor substrate recognition. For example, substituting Ile141 in the NTD of *Camellia sinensis* UGT94P1 for a serine resulted in a shift from xylosyltransferase to glucosyltransferase activity. This can be explained by the steric hindrance of the bulky side chain on isoleucine that only allows pentoses without C6 functional groups to fit into the catalytic pocket, while the presence of serine with a smaller side chain enables the accommodation of glucose and hydrogen bonds with C6-OH of glucose leading to a tuned glycosyltransferase activity ^41^. Overall, the CTDs mainly contribute to UDP-sugar interactions and recognitions by UGTs, and residues from both domains that affect intermolecular interactions with sugar moieties of UDP-sugars in the catalytic pocket should also be taken into account when modifying sugar donor specificities through rational protein engineering. The information provided by our research could yield more understanding of the biochemical and structural features of GT61, and thus help facilitate plant cell wall modifications to generate better feedstocks for biofuel production.

## Methods

### Protein expression and purification of SbXAT

#### Expression

The nucleotide sequences of *SbXAT* and *SbXXT* were codon-optimized for the mammalian expression system and synthesized (Biomatik Corporation). Gene fragments for different truncated versions of SbXAT (aa94-685, aa198-685 and aa286-685) and SbXXT (aa93-554 and 117-554) were amplified through PCR and cloned into the pDONR™221 vector using Gateway® BP Clonase™ II Enzyme mix (Thermo Fisher Scientific), followed by recombination with pGEn2-DEST expression vector using Gateway® LR Clonase™ II Enzyme mix (Thermo Fisher Scientific) as described previously ^42^. The obtained expression constructs coding for the GT61-GFP fusion proteins were transfected into 3 x 200 mL HEK-293 GnT1-cells at a density of 2.4E6 cell/ml using 3 µg/ml plasmid and 9 µg/ml polyethylenimine (PEI). The culture supernatants were harvested 5 days post transfection by centrifuging the culture fluid at 1000 x *g* for 10 min.

#### Purification

The clarified culture supernatant had imidazole added to 20 mM prior to loading onto a 5 ml HisTrap column on an AKTA PURE FPLC. Buffer A was 25 mM HEPES, pH 7.4, 250 mM NaCl. Buffer B was Buffer A + 500 mM imidazole. The bound protein was eluted with a 25 column volume (CV) gradient from 4-65% buffer B. The best elution fractions were pooled, dialyzed into 25 mM Tris, pH 8.0, 250 mM NaCl and treated with tobacco etch virus (TEV) to cleave off GFP and EndoF to remove the glycosylation moieties. The TEV and EndoF treated protein was put through a 5 mL HisTrap column to separate the GT61 catalytic domains from GFP, un-cleaved fusion protein, and the TEV and EndoF enzymes. The flow through and wash from the HisTrap column were pooled and dialyzed to 25 mM Tris, pH 8.0, 50 mM NaCl. The dialyzed protein was loaded onto a 5mL Q column on the AKTA Pure. Buffer A was dialysis buffer and B was 25 mM HEPES, pH 7.4, 1 M NaCl. The protein was eluted using a linear gradient from 0-75% buffer B in 25 CV. The cleanest elution fractions were pooled, concentrated and then injected on a 10×300 S75 increase column that was equilibrated in 25 mM Tris, pH 8.0, 100 mM NaCl. The cleanest elution fractions were pooled and concentrated to 10.3 mg/ml.

### Crystallization, X-rays data collection and structure determination of SbXAT

Crystallization trials were set up with the protein at 10.3 mg/ml using the Art Robbins Phenix robot. 0.2 µl of protein and 0.2 µl well solution were used for each sitting drop. The SbXAT sample was screened against the crystallization set of solutions: Berkeley Screen ^43^, MCSG-1 (Anatrace), ShotGun (Molecular Dimensions), PEG/Ion, Index, Crystal Screen, and PEGRx (Hampton Research). Crystals of SbXAT were found in PEG/Ion Screen condition E8 composed of 0.2 M sodium malonate pH 7.0 and 20 % (w/v) PEG 3,350. The crystal of SbXAT was placed in a reservoir solution containing 20% (v/v) glycerol, then flash-cooled in liquid nitrogen. The X-ray data set for SbXAT was collected at Stanford Synchroton Radiation Lightsource (SSRL) beamline 12-1. The diffraction data was processed using the program Xia2 ^44^. The crystal structure of SbXAT was solved by molecular replacement with the program PHASER ^45^ using as an initial coordinate the SbXAT model generated by ALPHAFOLD ^18^. The atomic positions obtained from the molecular replacement were used to initiate refinement within the Phenix suite ^46^. Structure refinement was performed using the phenix.refine program ^47^. Manual rebuilding was done using COOT ^48^. Root-mean-square deviations from ideal geometries for bond lengths, bond angles and dihedral angles were calculated with Phenix.refine ^47^. The stereochemical quality of the final model of SbXAT was assessed by the program MOLPROBITY^49^. Summary of crystal parameters, data collection, and refinement statistics can be found in Supplementary Table S1.

### *In vitro* activity assays

UDP-β-L-Ara*f* was purchased from Biosynth International, Inc. (Louisville, KY, USA), UDP-α-D-Xyl was purchased from Carbosource Service (Complex Carbohydrate Research Center, Athens, GA, USA), UDP-α-D-Xyl disodium was purchased from MedChemExpress (Monmouth Junction, NJ, USA). Xylooligosaccharides were purchased from MegaZyme (Bray, Ireland). SbXAT aa94-685 was used in the initial characterization of enzyme functionality and acceptor specificity, while SbXAT aa198-685 was used in all other enzyme assays. SbXXT aa93-554 was used in all activity assays if not specifically indicated.

Generally, a 20 μl reaction containing 4 μM protein, 1 mM UDP-sugar, 0.2 mM oligosaccharide acceptor, 1.25 mM (SbXXT) or 2.5 mM (SbXAT) MnCl_2_ and MgCl_2_ in 50 mM HEPES-Na at pH 7.0 (SbXXT) or pH 7.2 (SbXAT) was incubated at room temperature for different periods of time as indicated. The pH and divalent cation concentrations were optimized for each enzyme in preliminary experiments. The reaction products were treated with ion exchange resin (Dowex 50W X8), and the supernatant was mixed (1:1) with 2,5-dihydroxybenzoic acid (20 mg/ml in 50% methanol) matrix and dried on a metal plate. The samples were then analyzed by matrix-assisted laser desorption/ionization time-of-flight mass spectrometry (MALDI-TOF MS) using a UltrafleXtreme spectrometer (Bruker). The positive ion spectra were generated by summation of 2000 laser shots, and the obtained data was analyzed through flexAnalysis software (Bruker). For analysis of substrate specificity of GT61s, similar reactions were carried out with enzymes (1 μM SbXAT or 4 μM SbXXT) incubated with 50 μM UDP-sugars and 40 μM of separate xylo-oligosaccharides (Xyl_2_, Xyl_3_, Xyl_4_, Xyl_5_, Xyl_6_, and XA^2^XX/XA^3^XX arabinosylated xylotetraose mixture) in a 25 μL reaction for 15 min (SbXAT) and 75 min (SbXXT) in the same buffer condition described above for each enzyme. The UDP by-products generated from the reactions were quantified by the UDP-Glo™ Glycosyltransferase Assay kit (Promega) by measurement of luminescence using a BioTek Synergy H1 microplate reader (BioTek® Instruments, Inc.). The values were calculated after subtraction of background values obtained from incubations without acceptors. To compare the enzymatic activities between wildtype GT61 enzymes and their variants, similar reactions were executed with enzymes (0.5 μM SbXAT or 1.5 μM SbXXT) incubated with 300 μM UDP-sugars and 100 μM Xyl_6_; the UDP-Glo™ Glycosyltransferase Assay kit was then used to quantify UDP by-products after overnight incubation. The control experiments contained inactive enzymes (95℃ treated for 10 min), and the obtained values were subtracted from the data of enzymatic reactions.

### Analysis of the dissociation constant (K_d_) by MicroScale Thermophoresis (MST)

To determine the binding affinities of SbXAT, SbXXT, and their variants for UDP-Ara*f* and UDP-Xyl, 200 nM proteins (without prior TEV and EndoF cleavage) were incubated with the UDP-sugars at 16 different final concentrations ranging from 0 mM to 10 mM that were prepared through a two-fold serial dilution as described previously ^50^. The buffer environment used for the binding assay is the same as the reaction buffer used for activity assays mentioned above with addition of 0.05% Tween-20 to prevent protein aggregation and adsorption. The obtained 16 samples were incubated at room temperature for at least 10 min before being loaded into the standard treated glass capillaries. For MST analysis, 20% excitation power at 460-490 nm wavelength through the Nano Blue channel and 40% laser power were applied to all experiments at 22°C on a NanoTemper® Monolith NT.115 (NanoTemper Technologies, Germany) accompanied with blue/red filter. Data analysis was performed using MO.Affinity Analysis software (version 2.3, NanoTemper Technologies) to calculate the dissociation constants (K_d_) and plot a dose-response curve as normalized fluorescence change in MST signal (Fnorm) against the ligand concentration.

### Analysis of enzyme kinetics

To analyze the kinetic parameters of GT61 proteins, enzymes (0.5 μM) were incubated with a constant concentration of UDP-Ara*f* (25 μM) and varied concentrations of Xyl_6_ from 0 to 4 mM in 50 mM HEPES-Na, pH 7.2, with MgCl_2_ (2.5 mM) and MnCl_2_ (2.5 mM) for 30 min at room temperature. Reactions were terminated by directly adding the UDP Detection Reagent prepared from the UDP-Glo™ Glycosyltransferase Assay kit (Promega) according to the manufacturer’s protocol followed by measurement of luminescence using a BioTek Synergy H1 microplate reader (BioTek® Instruments, Inc.). The UDP by-products generated in each enzymatic reaction were calculated after subtraction of the background value from control experiments without enzymes, and the obtained values were used to calculate enzymatic velocities. Kinetic analyses were performed using nonlinear regression parameters and Michaelis-Menten equation by GraphPad Prism version 10.0.2 (GraphPad Software, Inc.).

### Molecular docking

To understand the protein-ligand interactions between SbXAT and UDP-Ara*f*, we used AutoDock Vina ^51^ to dock UDP-Ara*f* to the active site of the crystal structure of SbXAT through UCSF Chimera ^52^. The 3D structure of UDP-Ara*f* was downloaded from the PubChem database (https://pubchem.ncbi.nlm.nih.gov; CID: 44517206) and minimized using Chimera to prepare for molecular docking. The grid box was set to cover the entire donor substrate-binding site according to the structure of the POMGNT2 co-complex (PDB: 7E9J), and the conserved residues hypothetically interacting with the diphosphate moiety of UDP (Asn394, Arg515, Arg519) were placed in the center of the grid box. Exhaustiveness of search was set to 8, and the maximum energy difference was set to 3 kcal/mol for the docking process. Water molecules and all non-standard residues in the crystal structure of SbXAT were ignored. Ten binding modes were generated from the results of molecular docking, and the mode with the lowest estimated binding energy (ΔG) was selected and used as the optimal docked model. The predicted structure of SbXXT was generated using the AlphaFold Protein Structure Database (https://alphafold.ebi.ac.uk) ^18,23^ with UniProt ID: C5X460, followed by structural alignment with the crystal structure of SbXAT.

### Site-directed mutagenesis

To substitute the key substrate-binding residues in the catalytic sites of GT61s, the Q5® Site-Directed Mutagenesis Kit (New England Biolabs, Inc.) was used with mutagenic primers designed to generate substitution-incorporated SbXAT-F596Y-pGEn2 and SbXXT-Y462F-pGEn2 expression plasmids according to the manufacturer’s protocol. Briefly, forward primers 5’-GGCTGGATAC<u>TAT</u>AGAGGCCCCT-3’ (for SbXAT F596Y) and 5’-CGGCGCTTAC<u>TTT</u>GGCGAGCCTG-3’ (for SbXXT Y462F) that contain the desired codon changes (underlined) were used together with reverse primers 5’-ACGAACTCCAGGCCGC-3’ (for SbXAT F596Y) and 5’-GCGGCCCAATCTGTG-3’ (for SbXXT Y462F) to amplify the full length of SbXAT-pGEn2 and SbXXT-pGEn2 plasmid templates. The PCR products with substituted nucleotides were treated with a Kinase-Ligase-DpnI (KLD) enzyme mix followed by transformation into NEB® 5-alpha Competent *E. coli* (New England Biolabs, Inc.). The plasmids prepared from the selected colonies were full-length sequenced (Plasmidsaurus, Inc.) and used for protein expression in HEK-293 cells as described above.

### Phylogenetic analysis

GT61 protein sequences in *Arabidopsis thaliana*, *Oryza sativa*, *Populus trichocarpa*, and *Sorghum bicolor* were retrieved from Phytozome v13 (https://phytozome-next.jgi.doe.gov) ^53^: Multiple GT61 protein sequences were aligned through Multiple Sequence Comparison by Log-Expectation (MUSCLE) algorithm ^54,55^ using SnapGene software (www.snapgene.com). The aligned sequences were then used to construct a phylogenetic tree based on the maximum likelihood method using MEGA11 ^56^ software, following by annotation and display through the online tool, iTOL v7.1 (https://itol.embl.de) ^57^.

## Data availability

The atomic coordinates and structure factors are available in the PDB under accession number 9OIS for SbXAT.

## Acknowledgements

This material is based upon work supported by the Joint BioEnergy Institute, U.S. Department of Energy, Office of Science, Biological and Environmental Research Program under Award Number DE-AC02-05CH11231 with Lawrence Berkeley National Laboratory. Use of the Stanford Synchrotron Radiation Lightsource, SLAC National Accelerator Laboratory, is supported by the U.S. Department of Energy, Office of Science, Office of Basic Energy Sciences under Contract No. DE-AC02-76SF00515. The SSRL Structural Molecular Biology Program is supported by the DOE Office of Biological and Environmental Research, and by the National Institutes of Health, National Institute of General Medical Sciences (P30GM133894). This work was also financially supported by the Center for Bioenergy Innovation (CBI), which is a US Department of Energy Bioenergy Research Center supported by the Office of Biological and Environmental Research Program in the DOE Office of Science under award number ERKP886.

## Author contributions

H.-T.W., H.V.S., P.D.A., and J.H.P. conceived and designed the project. A.M.DG., D.C. and K.W.M. expressed and A.M.DG., P.K.P. and B.R.U. purified proteins. H.-T.W. conducted sequence analyses, activity assays, molecular docking and mutagenesis. J.H.P. determined the crystal structure. H.-T.W., J.H.P. and H.V.S. wrote the manuscript with input from all authors.

---

**Table S1.** Statistics for data collection and refinement of SbXAT.

|  | SbXAT |
| --- | --- |
| **Data collection** |  |
| Wavelength (Å) | 0.97946 |
| Resolution range (Å) | 37.57 – 1.99 (2.06 - 1.99) |
| Detector Distance (mm) | 300 |
| Φ (deg.) collected / ΔΦ (deg.) | 200/0.25 |
| Exposure time (seconds) | 0.25 |
| Temperature of collect (Kelvin) | 100 |
| **Data statistics** |  |
| Space group | P 61 2 2 |
| Unit-Cell parameters (Å) | a=88.159 b=88.159 and c=211.509 |
| Total reflections | 68422 (6674) |
| Unique reflections | 34212 (3337) |
| Multiplicity | 2.0 (2.0) |
| Data completeness (%) | 99.87 (100.00) |
| <I/σ(I)> | 13.04 (1.95) |
| R<sub>merge</sub> (%) | 0.02524 (0.331) |
| R<sub>pim</sub> (%) | 0.02524 (0.331) |
| CC<sub>1/2</sub> | 0.999 (0.795) |
| **Structure Refinement** |  |
| Reflections used in refinement | 34205 (3337) |
| Reflections used for R<sub>free</sub> | 1677 (156) |
| R<sub>factor</sub> | 0.1990 (0.2947) |
| R<sub>free</sub> | 0.2284 (0.3318) |
| CC (work) | 0.960 (0.834) |
| CC (free) | 0.945 (0.842) |
| RMS from ideal geometry |  |
| Bond lengths (Å) | 0.006 |
| Bond angles (°) | 0.70 |
| Average B-factor | 50.04 |
| Macromolecules | 50.19 |
| Ligands | 45.09 |
| Solvent | 48.03 |
| Ramachandran Plot |  |
| Favored region (%) | 96.04 |
| Allowed region (%) | 3.69 |
| Outliers region (%) | 0.26 |
| Rotamer outliers (%) | 0.31 |
| Number of TLS groups | 8 |

**Fig. S1.**
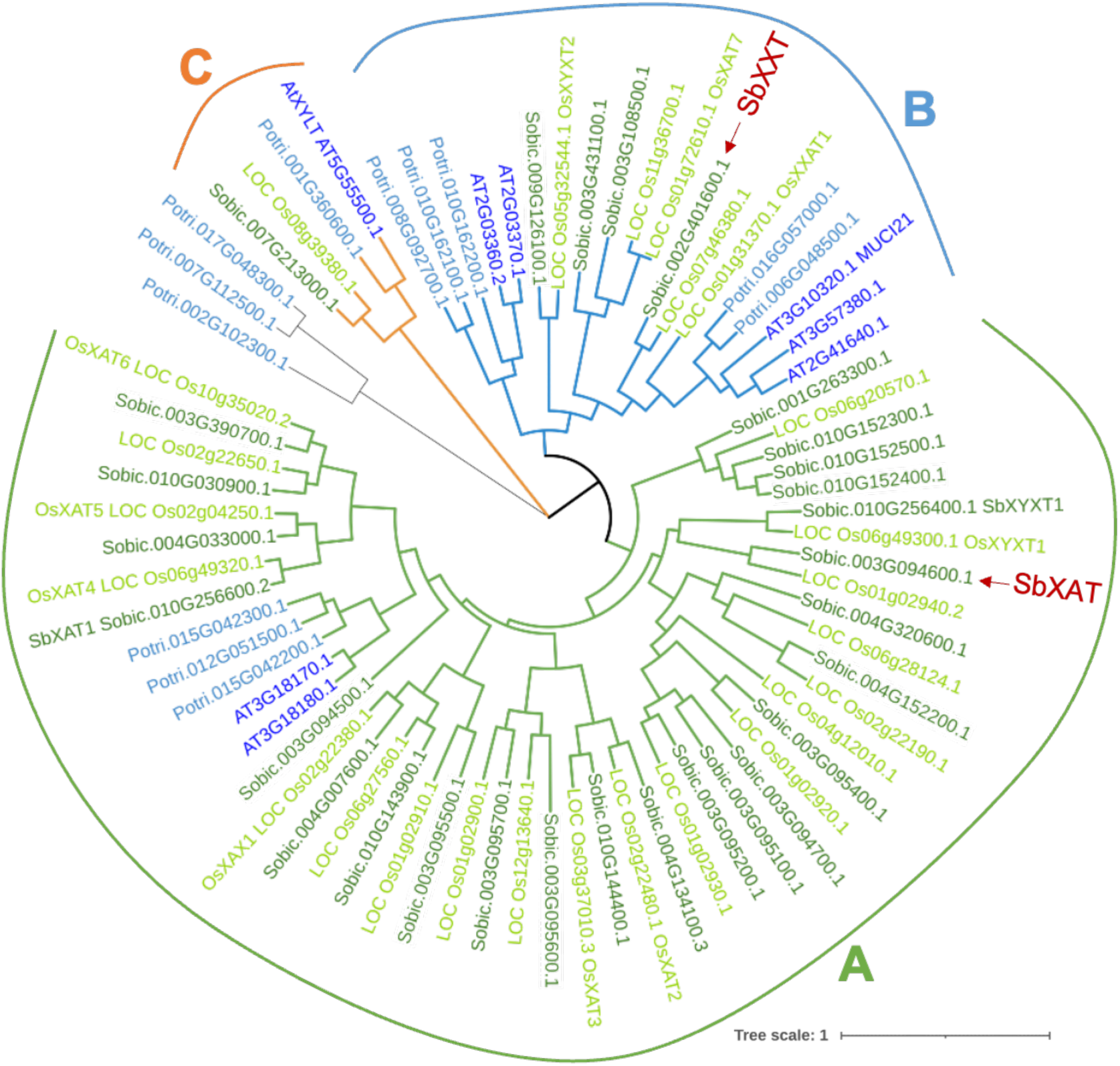
Phylogenetic tree of GT61 members from *Sorghum bicolor* (dark green), *Oryza sativa* (light green), *Arabidopsis thaliana* (dark blue), and *Populus trichocarpa* (light blue). The two sorghum GT61 proteins characterized in this study are indicated by red arrows, and clades are labeled A-C according to Anders et al. ^10^. The phylogenetic tree was constructed using maximum likelihood method.

**Fig. S2.**
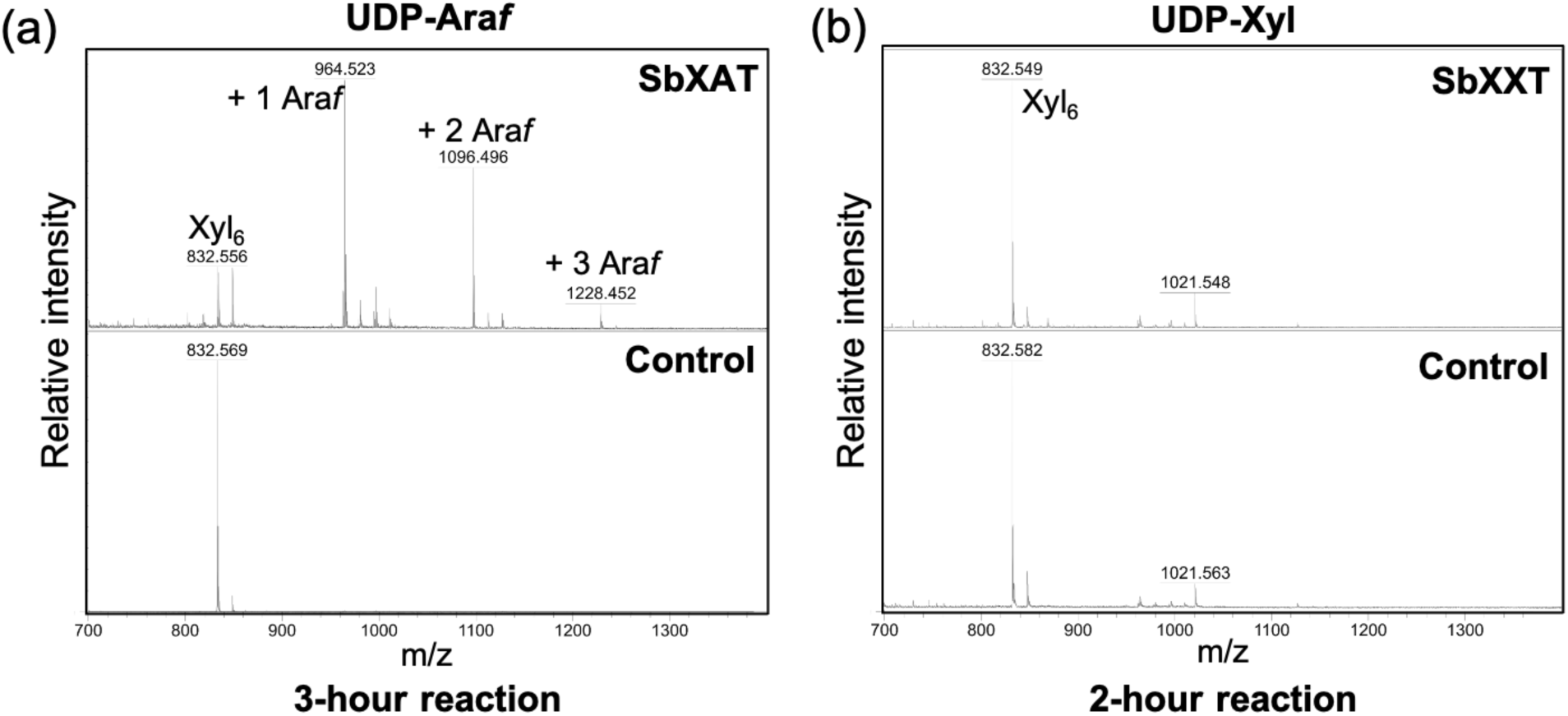
MALDI-TOF MS analysis of the products generated by SbXAT (a) and SbXXT(b) using UDP-Ara*f* and UDP-Xyl, respectively, as the donor substrates. Xyl_6_ was used as the acceptor in *in vitro* assays. The reaction time for each assay is indicated. Each transfer of an Ara*f* or Xyl increases the mass of the Xyl_6_ acceptor by 132 Da as shown by annotated [M+Na]^+^ ions. Results of overnight incubation are shown in Fig. 1.

**Fig. S3.**
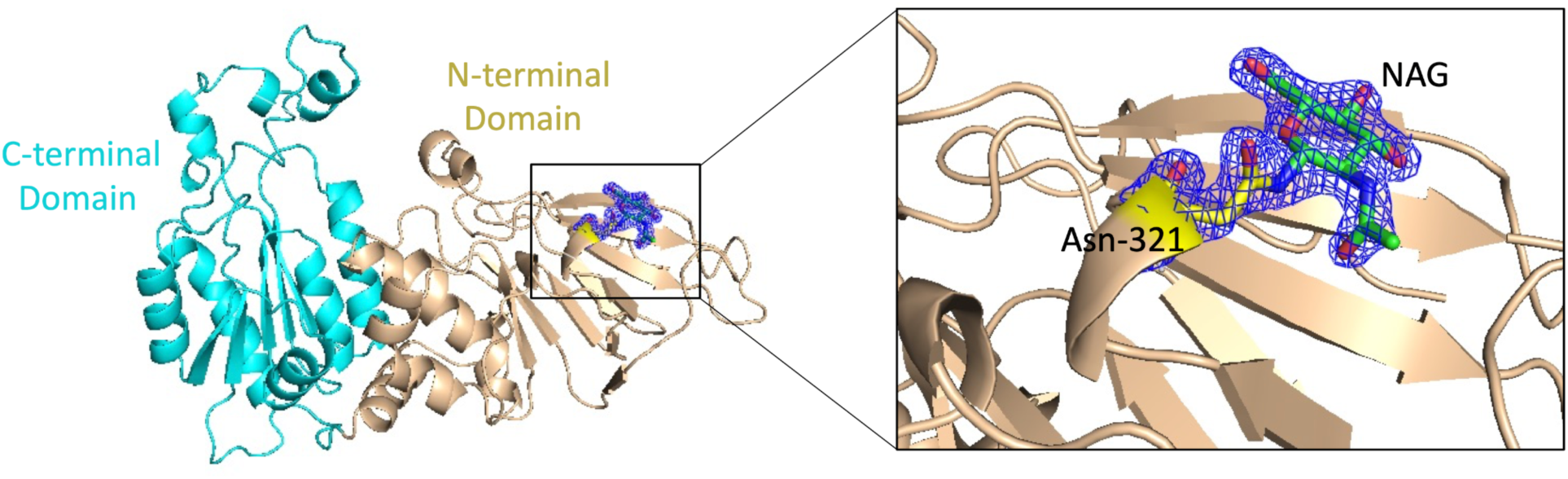
Glycosylation site observed at Asn321 in the SbXAT crystal structure. The electron density map around Asn-321 and the glycosylation sugar 2-acetamido-2-deoxy-beta-D-glucopyranose (NAG) was calculated using a 2mFo-DFc map contoured at 2.0 sigma level.

**Fig. S4.**
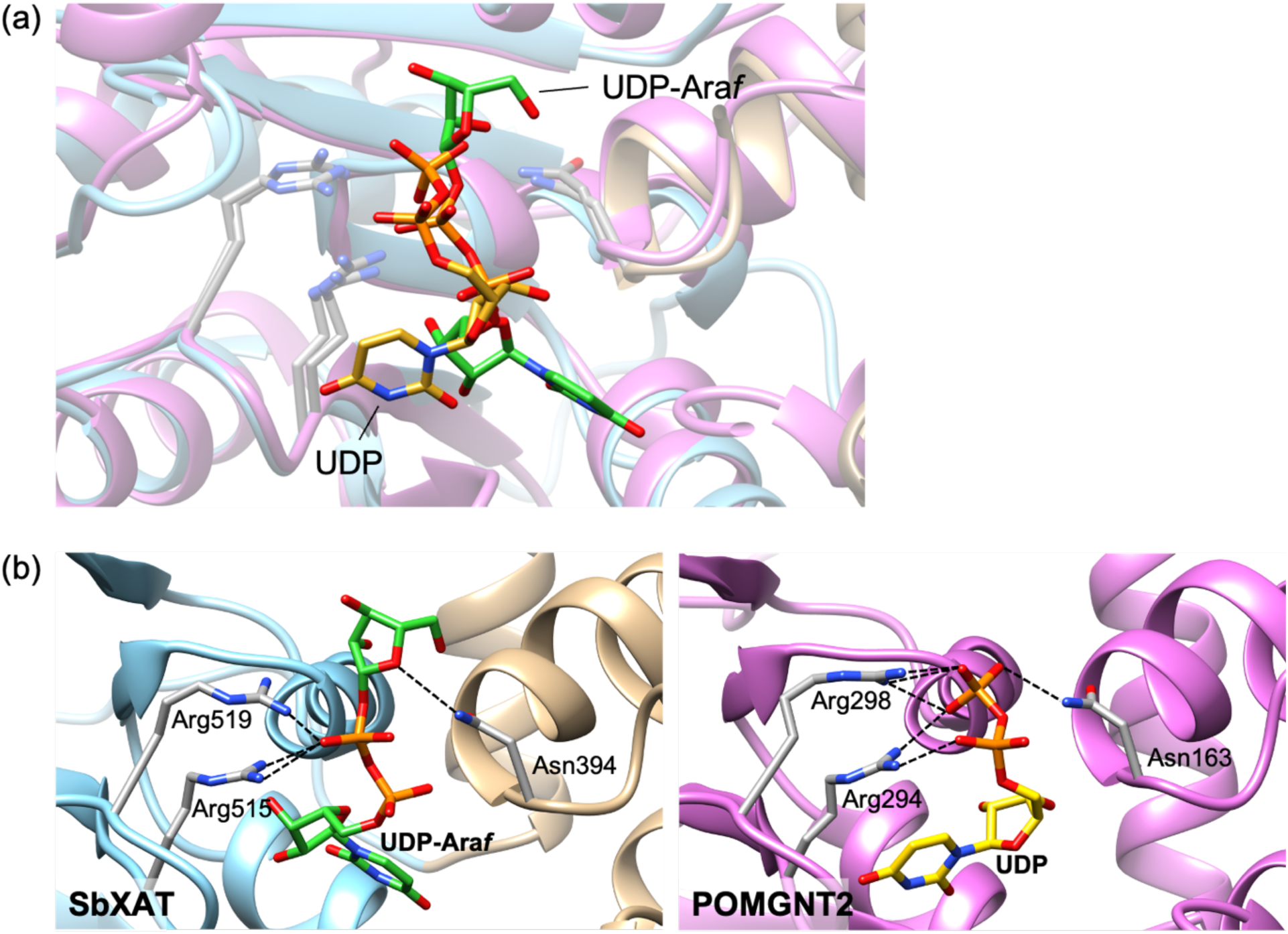
Comparison between the orientations of the docked UDP-Ara*f* and co-crystalized UDP in the active site of SbXAT and POMGNT2, respectively. **a,** Superposition of SbXAT (ribbons, N-lobe in tan, C-lobe in blue) in complex with the docked UDP-Ara*f* (green sticks) against the POMGNT2 structure (magenta ribbons) in complex with UDP (gold sticks) (PDB: 7E9J, chain A). **b**, Left: interactions between the docked UDP-Ara*f* (green sticks) and SbXAT (ribbons, N-lobe in tan, C-lobe in blue). Right: interactions between co-crystalized UDP (gold sticks) and POMGNT2 (magenta ribbons) (PDB: 7E9J, chain A). Hydrogen bonds are represented by dashed lines (black), and the conserved diphosphate-interacting residues in both structures are shown as gray sticks with annotations.

**Fig. S5.**
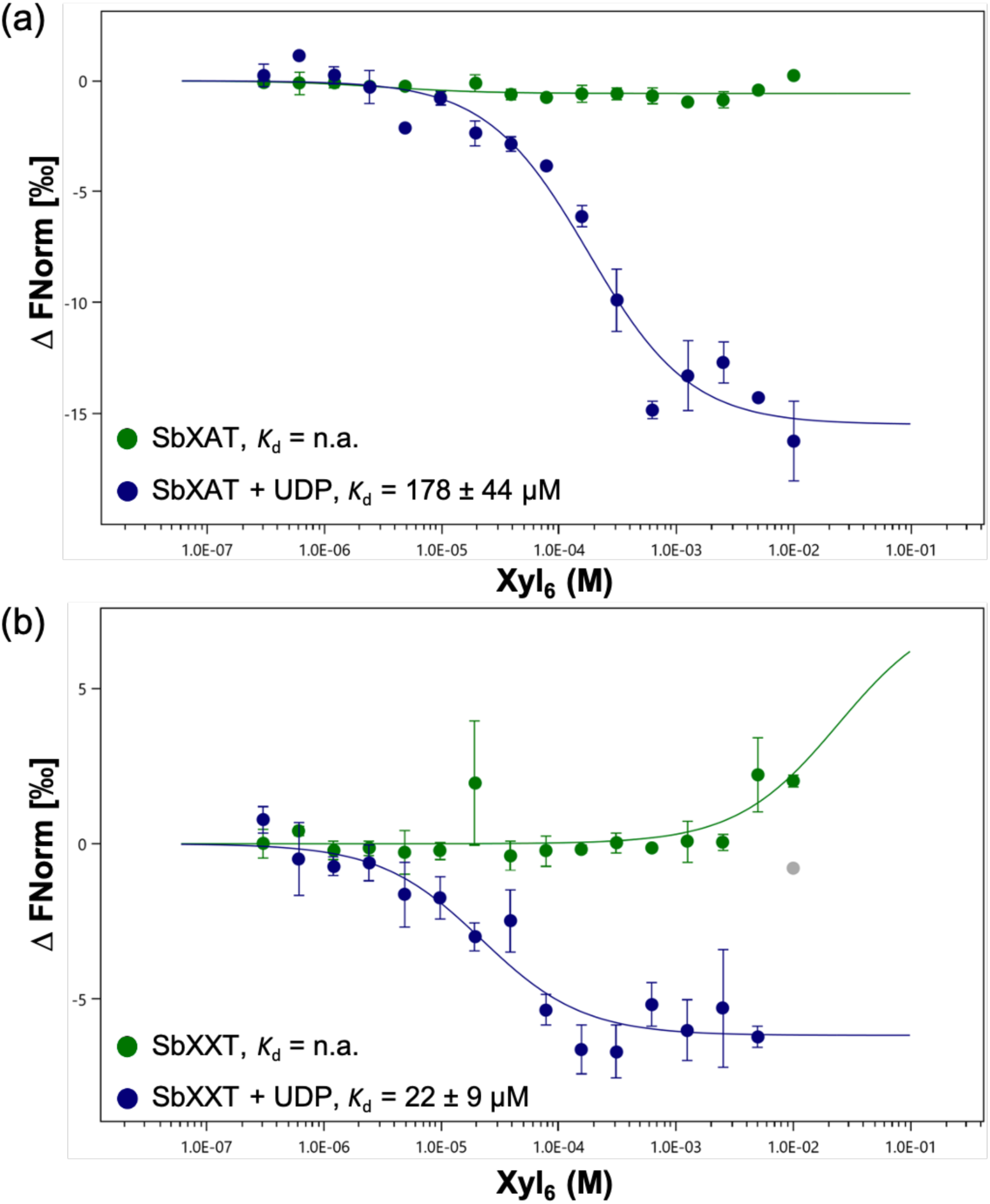
Substrate-binding affinity analysis of SbXAT and SbXXT for Xyl_6_ acceptor substrate in the presence or absence of UDP through microscale thermophoresis (MST). Normalized binding curves of SbXAT (**a**) and SbXXT (**b**) for Xyl_6_ with (blue) and without (green) UDP. In the absence of UDP, no binding was observed. The dissociation constants (*K*_d_) ± SD in the presence of UDP are indicated in the figure. The concentration of SbXAT or SbXXT was kept constant (200 nM), while Xyl_6_ concentration was varied from 0 to 10 mM. In assays in the presence of UDP, a concentration of 1 mM was used. Data points in gray were ignored due to protein aggregation. Error bars represent means ± SD from three technical replicates (n = 3).

**Fig. S6.**
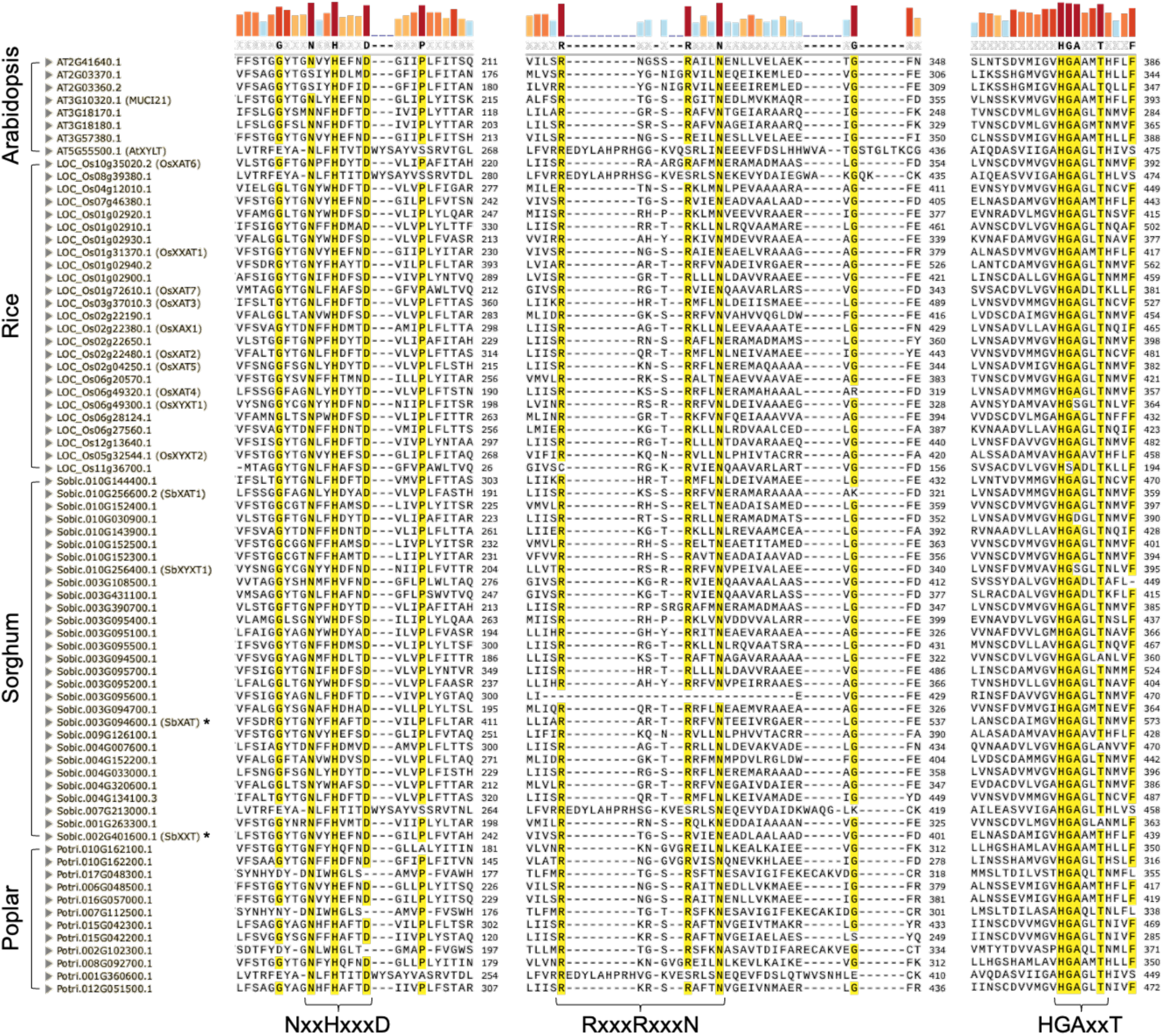
Protein sequence alignment of GT61 members from *Arabidopsis thaliana*, rice (*Oryza sativa*), sorghum (*Sorghum bicolor*), and poplar (*Populus trichocarpa*). The highly conserved motifs NxxHxxxD, RxxxRxxxN, and HGAxxT are indicated. Colored bars on the top represent the sequence consensus, and residues > 90% consensus threshold are highlighted in yellow. MUSCLE was used for multiple sequence alignment. *, the two sorghum GT61s identified in our study.

**Fig. S7.**
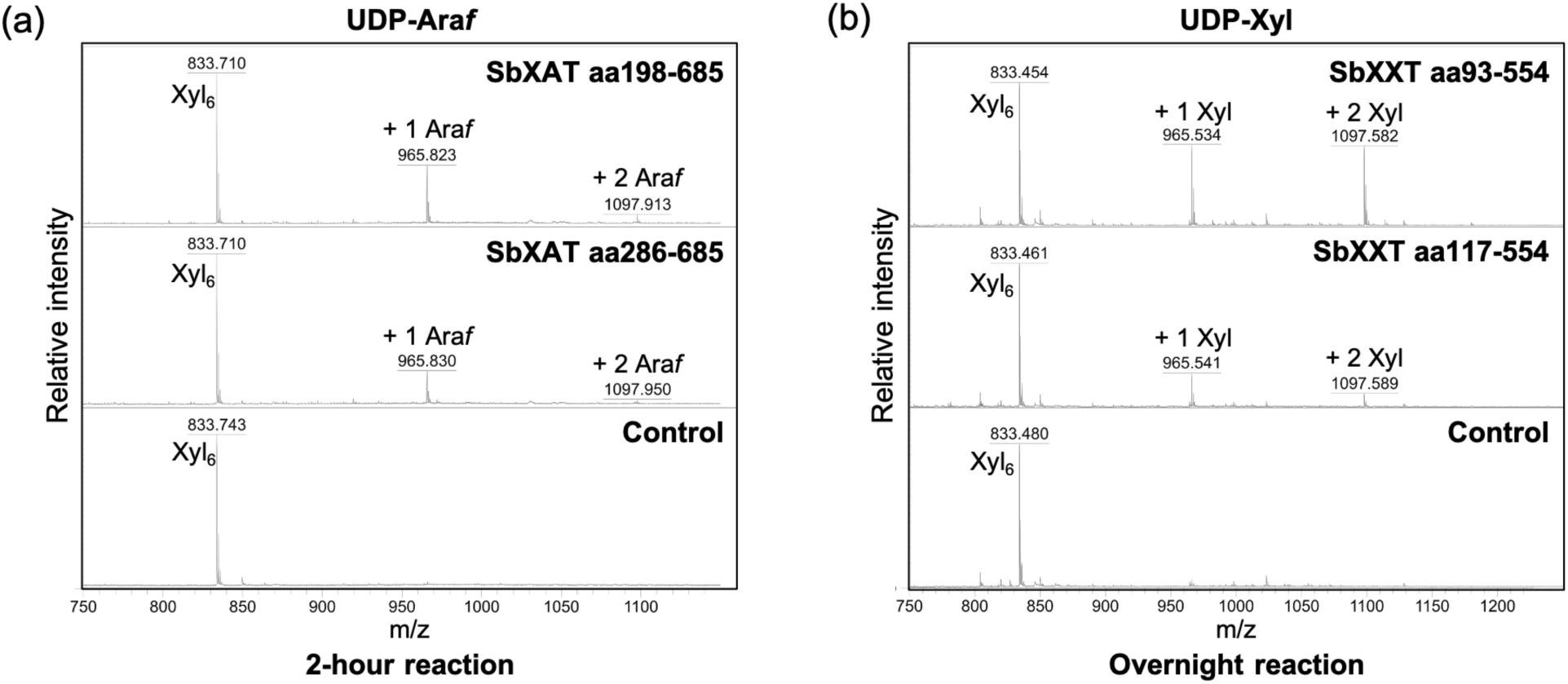
Different glycosyltransferase activities of the N-terminal flexible region-truncated GT61s as revealed by MALDI-TOF MS analysis. Products generated by SbXAT (**a**) and SbXXT (**b**) using UDP-Ara*f* and UDP-Xyl, respectively, as the donor substrates and Xyl_6_ as the acceptor in the *in vitro* assays were analyzed. The reaction time and the length of enzymes used for each assay are indicated. Each transfer of an Ara*f* or Xyl increases the mass of the Xyl_6_ acceptor by 132 Da as shown by annotated [M+Na]^+^ ions. The results show that truncation variants that lack part of the N-terminal disordered region are less active.

